# Sperm exposure to accessory gland secretions alters the transcriptomic response of the endometrium in cattle

**DOI:** 10.1101/2022.01.06.475202

**Authors:** José María Sánchez, María Belén Rabaglino, Sandra Bagés-Arnal, Michael McDonald, Susanta K Behura, Thomas E Spencer, Pat Lonergan, Beatriz Fernandez-Fuertes

**Author notes:** **Corresponding Author:** Pat Lonergan, School of Agriculture and Food Science, University College Dublin, Belfield, Dublin 4, Ireland, /6012147, Beatriz Fernandez-Fuertes, Department of Animal Reproduction, National Institute for Agriculture and Food Research and Technology (INIA), Spanish National Research Council (CSIC), Av. Puerta de Hierro, 18, 28040 Madrid, Spain.

## Abstract

In a recent study from our group, mating to intact, but not vasectomised, bulls modified the endometrial transcriptome, suggesting an important role of sperm in the modulation of the uterine environment in this species. However, it is not clear whether these changes are driven by intrinsic sperm factors, or by factors of accessory gland (AG) origin that bind to sperm at ejaculation. Thus, the aim of the present study was to determine whether ejaculated sperm, which are suspended in the secretions of the AGs, elicit a different endometrial transcriptomic response than epididymal sperm, which have never been exposed to AG factors. To this end, bovine endometrial explants collected from heifers in oestrus were incubated alone (control), or with epididymal or ejaculated sperm. RNA-sequencing revealed 1912 differentially expressed genes (DEGs) between in endometrial explants exposed to epididymal sperm compared with control explants, whereas 115 DEGs genes detected between endometrial explants exposed to ejaculated sperm in comparison to control explants. In both cases, the top pathways associated with these genes included T cell regulation and NF-KB and IL17 signalling. To confirm whether AG factors were directly responsible for the dampening of the endometrial response elicited by ejaculated sperm, endometrial explants were incubated with epididymal sperm previously exposed, or not, to seminal plasma (SP). Exposure to SP abrogated the downregulation of *SQSTM1* by epididymal sperm, and partially inhibited the upregulation of *MYL4* and *CHRM3* and downregulation of *SCRIB*. These data indicate that factors of AG origin modulate the interaction between sperm and the endometrium in cattle.

## INTRODUCTION

During their transit through the male reproductive tract, sperm are sequentially bathed in secretions derived from the rete testes, epididymides, and accessory sex glands (AGs), which collectively constitute a complex fluid termed seminal plasma (SP). The importance of epididymal fluid for sperm function is clear, as progressive acquisition of motility^1^, the ability to bind to the oviductal epithelia^2^, and the ability to interact with oocyte vestments^1^ takes place as sperm move from the caput epididymis to the cauda, where they are stored prior to ejaculation. As evidence of the complexity of the events that take place in this organ, sperm epididymal maturation takes between 1-2 weeks to complete in most species^5^. In contrast, considering that bovine sperm can be found in the oviduct minutes after artificial insemination (AI) or mating^6^, the duration of sperm exposure to the secretions of the AGs at ejaculation is minimal. Despite this, proteins secreted by AGs can bind to the sperm plasma membrane^7^, and seem to affect sperm function as evidenced by an increase in progressive motility^8^, in the ability to bind to oviductal cells^9^, and in the ability to bind to the zona pellucida^10^, when cauda epididymal sperm are exposed to SP.

In addition to affecting sperm function, SP has been proposed to regulate the maternal environment. In mammals, including humans^11^, mice^12–14^, and livestock species such as horses^15^, pigs^16^, and sheep^17^, exposure to semen induces an inflammatory response in the female reproductive tract. It has been suggested that SP components are responsible for this modulation of the maternal immune response, creating an immune tolerogenic environment^18,19^ which can lead to improved embryo development and fertility. Indeed, in mice, removal of the seminal vesicles (the main AG contributing to SP volume in that species) leads to impaired embryo development, a reduction in the number of implantation sites and, ultimately, decreased pregnancy rates^20^. Similarly, infusion of SP into the uteri of sows at the time of AI can impact embryo development^16,21^, whereas in mares, SP has been shown to improve fertility after AI into a uterus in which endometritis was previously induced^22^.

In cattle, however, the role of SP in fertility is not clear. Despite inducing changes in the expression of endometrial cytokines^23^, uterine infusion of SP at the time of AI^24,25^ or exposure of females to vasectomized bulls for 21 days prior to AI^26^ failed to improve pregnancy rates. Importantly, in contrast to the mouse, horse, and pig, where the ejaculate comes into direct contact with the endometrium^27–29^, bulls ejaculate in the anterior vagina^30^, and it is questionable whether any of the fluid fraction reaches the uterus. In fact, SP infusion into the vagina, but not into the uterus, can modify endometrial epidermal growth factor concentrations in repeat breeder cows^31^. We recently reported a modest increase in Day 14 conceptus length following transfer of bovine embryos on Day 7 into the uterus of heifers mated to a vasectomized bull in comparison to unmated heifers^32^. In addition, data from our group have shown that mating to intact or vasectomised bulls induces a higher expression of interleukin 1 A (*IL1A*) and tumor necrosis factor alpha (*TNFA*) in the vagina than in the oviduct, whereas this difference is not observed in unmated heifers^33^. However, mating to intact bulls, but not to vasectomised bulls, induces modest changes in the endometrial transcriptome 24 h after mating^33^. These data suggest that, in species that ejaculate intravaginally, sperm might play a more critical role in modulating the endometrial environment, perhaps acting as a vehicle for factors derived from the AGs. An example of such species-specific differences regarding the modulation of the uterine environment by sperm or SP, is the regulation of *IL17*. In mice, *Il17* expression increased after SP exposure, regardless of the presence or absence of sperm^14^. In contrast, in cattle, upregulation of *IL17* has been observed after mating to intact, but not vasectomised bulls^33^, as well as after AI with semen (SP and sperm) but not SP alone^23^. Moreover, bull sperm have been reported to induce a pro-inflammatory response in epithelial uterine cells *in* vitro^34,35^. However, it is not clear whether these effects are triggered by AG factors incorporated onto the sperm surface at the time of ejaculation, or by molecules acquired during spermatogenesis and epididymal maturation.

Since sperm in the cauda epididymis have never been in contact with AG secretions, comparing the response induced in the endometrium by cauda epididymal sperm with that induced by ejaculated sperm is a suitable model to evaluate the potential effect of AG-derived factors. Based on the evidence in other species supporting a beneficial effect of SP, we hypothesised that ejaculated sperm, which have been exposed to SP at the time of ejaculation, induce a different modulation of endometrial gene expression than epididymal sperm. To test this hypothesis, using a previously validated endometrial explant model^36–38^, we compared the transcriptomic changes elicited in bovine endometrial explants following co-culture with epididymal or ejaculated sperm. Because exposure to epididymal sperm resulted in a much higher number of differentially expressed genes (DEGs) than incubation with ejaculated sperm, we further investigated the modulation of the sperm-endometrial interaction by SP by incubating explants with epididymal sperm, previously exposed or not to SP.

## MATERIAL AND METHODS

All experimental procedures involving animals were approved by the Animal Research Ethics Committee of University College Dublin and licensed by the Health Products Regulatory Authority, Ireland, in accordance with Statutory Instrument No. 543 of 2012 (under Directive 2010/63/EU on the Protection of Animals used for Scientific Purposes).

Unless otherwise stated, all chemicals and reagents were sourced from Sigma-Aldrich (Arklow, Ireland).

### Experimental design

Endometrial explants were generated from the uteri of oestrous synchronised heifers recovered 24 h after the onset of oestrus, as detailed in the following sections. The experimental design is summarized in Fig. 1.

**Fig. 1.**
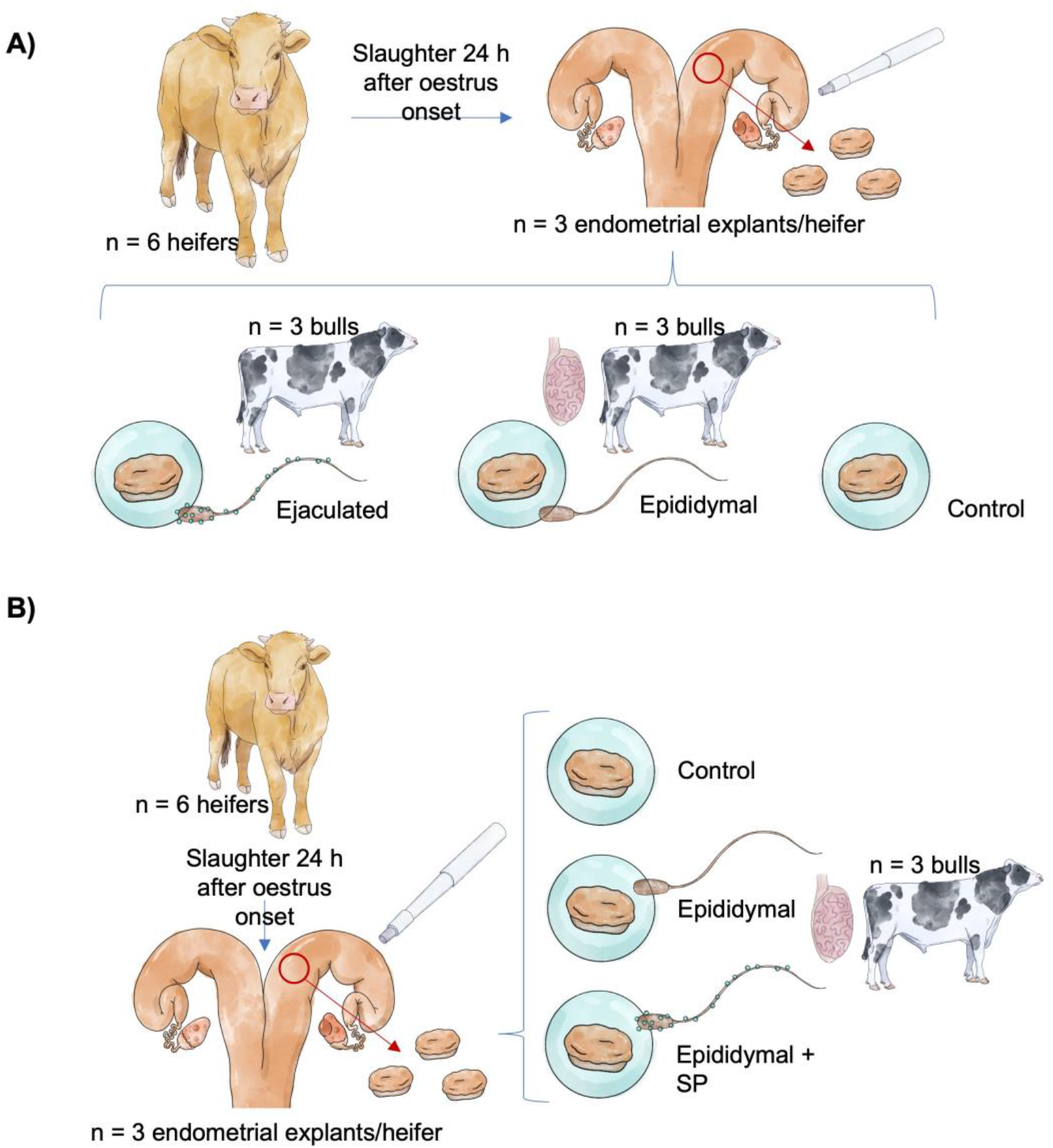
**A)** Heifers (n = 6) were oestrus synchronised and slaughtered approximately 24 h after being observed in standing oestrus. Endometrial explants (n = 3 per animal) were recovered from the ipsilateral horn and incubated for 6 h with: 1) RPMI medium (Control), 2) RPMI medium + 5 × 10^6^ epididymal sperm (Epididymal), and 3) RPMI medium + 5 × 10^6^ ejaculated sperm (Ejaculated). Subsequent RNA sequencing of explants was conducted. **B)** Explants recovered from oestrus synchronised heifers slaughtered approximately 24 h after oestrus onset (n = 6 heifers, n= 3 explants per animal), were incubated for 6 h with: 1) Roswell Park Memorial Institute-1640 (RPMI) medium (Control), 2) RPMI medium + 5 × 10^6^ epididymal sperm (Epididymal), and 3) RPMI medium + 5 × 10^6^ epididymal sperm previously exposed to seminal plasma (Epididymal + SP). Expression of a set of candidate genes was then analysed by qPCR.

The aim of Experiment 1 was to determine whether ejaculated sperm, which have been exposed to SP at the time of ejaculation, induce a different modulation of endometrial gene expression than epididymal sperm (which have had no contact with AG secretions). To that end, RNA-sequencing was performed on endometrial explants that had been incubated for 6 h with 1) Roswell Park Memorial Institute-1640 (RPMI) medium alone (Control), 2) RPMI medium + 5 × 10^6^ epididymal sperm (Epididymal), or 3) RPMI medium + 5 × 10^6^ ejaculated sperm (Ejaculated; Fig. 1A).

Because the number of DEGs detected between control explants and explants exposed to ejaculated sperm was so small, in comparison to the number of DEG between control and epididymal sperm explants, it was hypothesised that exposure of epididymal sperm to SP in vitro would modulate the endometrial response. To test this hypothesis, in Experiment 2, endometrial explants were incubated for 6 h with 1) RPMI medium alone (Control), 2) RPMI medium + 5 × 10^6^ epididymal sperm (Epididymal), or 3) RPMI medium + 5 × 10^6^ epididymal sperm previously incubated with SP (Epididymal + SP; Fig 1B). Relative expression of a set of candidate genes (selected based on the fold-change difference observed in Experiment 1 or their ability to serve as biomarkers of treatment with epididymal sperm, was analysed by quantitative real-time PCR (qPCR).

### Generation of endometrial explants

Crossbreed beef heifers (n= 6 for Experiment 1, and n= 6 for Experiment 2) were oestrous synchronised using an 8-day intravaginal device (PRID® Delta, 1.55 g progesterone, Ceva Santé Animale, Libourne, France), together with a 2 mL intramuscular injection of a synthetic gonadotrophin releasing hormone (Ovarelin®, equivalent to 100 μg Gonadorelin, Ceva Santé Animale) administered on the day of PRID insertion. One day prior to PRID removal, all heifers received a 5 mL intramuscular injection of prostaglandin F2 alpha (Enzaprost®, equivalent to 25 mg of Dinoprost, Ceva Santé Animale) to induce luteolysis. Heifers were slaughtered approximately 24 h after being observed in standing oestrus, and their reproductive tracts were recovered and transported on ice to the laboratory. All heifers exhibited a large preovulatory follicle (15.5 ± 0.9 mm; mean diameter ± standard error of the mean, SEM; Supp. Table 1).

Uteri were processed as described previously Mathew et al.^36^. Briefly, the uterine horn ipsilateral to the pre-ovulatory follicle was opened longitudinally with sterile scissors, and the exposed endometrium was washed with Dulbecco’s phosphate-buffered saline solution (PBS) supplemented with 1% Antibiotic-Antimycotic (ABAM; Gibco, ThermoFisher Scientific, Dublin, Ireland). Three tissue samples from each animal were obtained from intercaruncular areas from the middle of the uterine horn with the use of a sterile 8 mm biopsy punch (Stiefel Laboratories Ltd, High Wycome, UK). Sterile blades were used to dissect the endometrium away from the myometrium. Once dissected, the explants were washed twice in Hank’s balanced salt solution (Gibco, ThermoFisher Scientific) containing 1% ABAM, and then transferred into a 24-well plate, so that each well contained one explant in 1 ml RPMI medium (11835-030, Gibco, ThermoFisher Scientific) supplemented with 1% ABAM. Explants were cultured endometrial side up at 39 °C under an atmosphere of 5% CO_2_ for 4 h before use.

After 4 h equilibration in RPMI medium, explants were transferred to wells containing the different treatments described above and incubated for 6 h at 39 °C under an atmosphere of 5% CO_2_. After incubation, they were washed twice in RPMI media to remove sperm, snap frozen, and stored at −80 °C until further analysis. Six replicates were carried out for each experiment, each replicate corresponding to a synchronised heifer. To minimise variation, in each experiment, explants from the same uterus were used across all treatments in each replicate.

### Preparation of sperm and SP

Ejaculated sperm was obtained from 3 Holstein Friesian bulls collected at a local AI centre (National Cattle Breeding Centre, Naas, County Kildare, Ireland) using an artificial vagina. Because bull SP contains high concentrations of RNase, which degrades endometrial-explant RNA in settings similar to those in the current study^39^, sperm were isolated from SP by washing through a density gradient. Thus, after pooling the ejaculates, 1 ml of sample was washed through a 90–45% discontinuous Percoll gradient (Pharmacia, Uppsala, Sweden) for 9 min at 700 g, followed by a second centrifugation in RPMI at 200 g for 5 min, to isolate ejaculated sperm from SP. Seminal plasma was obtained by centrifuging the rest of the pooled ejaculates for 9 min at 700 g, and filtering the supernatant through a 0.2 μm pore filter (Sarstedt, Wexford, Ireland).

To obtain cauda epididymal sperm, testes from 3 mature bulls were collected from a local abattoir and transported within 2 h to the laboratory at ambient temperature. A small incision was made in the cauda epididymis and the lumen of the deferent duct was cannulated with a blunted 22 G needle. Sperm were then gently flushed through the cauda with a 5 ml syringe loaded with PBS at 38 °C, as described previously Fernandez-Fuertes et al.^40^. Epididymal sperm recovered from the 3 bulls were pooled and washed through Percoll and RPMI as described above. In Experiment 2, prior to the Percoll wash, two aliquots of the epididymal sperm pool were centrifuged at 200 g for 5 min. Then, one aliquot was resuspended in 1 ml SP and the other in 1 ml RPMI. After a 5 min incubation, samples were washed through Percoll and RPMI.

In all instances, sperm concentration was assessed with a haemocytometer and adjusted to 5 × 10^6^ total sperm per ml per well, which is a typical concentration dose per straw of fresh semen used for AI^41^.

### Explant RNA extraction

Briefly, for total mRNA extraction, samples were first homogenized in Trizol reagent (Invitrogen, Carlsbad, CA) using a steel bead and the Qiagen tissue lyzer (2 × 120 s at 30 Hz). On-column RNA purification was performed using the Qiagen RNeasy kit (Qiagen, Crawley, Sussex, UK) per the manufacturer’s instructions. The quantity of RNA was determined using the Nano Drop 1000 spectrophotometer (Thermo Fisher Scientific).

### Experiment 1

#### RNA-sequencing

Prior to RNA sequencing analysis, the RNA quality was assessed by the Agilent Bioanalyzer (Agilent Technologies, Cork, Ireland). All samples originating from one heifer were discarded due to low RNA integrity number (RIN).

RNA library preparation and sequencing were performed by the University of Missouri DNA Core Facility as described previously Moraes et al.^42^. The raw sequences (fastq) were subjected to quality trimming control using fqtrim (https://ccb.jhu.edu/software/fqtrim/). Then, the quality reads were mapped to the bovine reference genome UMD3.1 using Hisat2 mapper (https://ccb.jhu.edu/software/hisat2/), which is a fast and sensitive alignment program of next-generation sequencing data^43^. Read counts mapping to each gene were determined from the mapping data using FeatureCounts^44^.

#### Bioinformatics

Raw counts were analysed with packages in the R software. As a first step, the distribution of the samples was assessed through a principal component analysis (PCA) to determine if the transcriptome was influenced by the heifer from which the endometrial sample was obtained. As the PCA plot revealed a heifer effect, this was removed with the CombatSeq function from the sva packages^45^ (Supp. Fig. 1). Subsequent analyses were performed with the DESeq2 package^46^.First, the read counts were normalized by library size with DESeq2 methods. Then, the read counts were modeled through a negative binomial distribution with a fitted mean and a gene specific dispersion parameter. Finally, the Wald test statistic was employed to test for model significance. P-values were adjusted with the Benjamini-Hochberg method, and the DEG were considered at False Discovery Rate (FDR) <0.05.

For the functional analysis, DEGs were interrogated for enriched KEGG pathways/Biological Processes (FDR <0.05) using the R package clusterProfiler^47^. The R package org.Bt.eg.db was used for genome-wide annotation for the bovine.

### Experiment 2

#### cDNA synthesis and qPCR analysis

For each sample, cDNA was prepared from 500 ng of total RNA using the High-Capacity cDNA Reverse Transcription Kit (Thermo Fisher Scientific) according to the manufacturer’s instructions. The purified cDNA was then diluted in RNase-and DNase-free water up to a volume of 300 μL and stored at -20°C for subsequent analysis.

All primers used to investigate changes in endometrial gene expression were designed using Primer Blast software (https://www.ncbi.nlm.nih.gov/tools/primer-blast/) (Supp. Table 1). Duplicate qPCR assays were performed in a total volume of 20 μL, containing 10 μL FastStart Universal SYBR Green Master (Roche Diagnostics Ltd., West Sussex, UK), 1.2 μL forward and reverse primer mix (300 nM final concentration), 2.6 μL nuclease-free water and 5 μL cDNA template on the ABI Prism 7500 Fast Sequence Detection System (Life Technologies). Thermo-cycling conditions were as follows: 95°C for 10 min for one cycle, followed by 95°C for 15 s, and 60°C for 1 min for 40 cycles. A dissociation curve was also added to ensure specificity of amplification. The presence of a single sharp peak in the melt curve analysis confirmed the specificity of all targets. A total of eight potential reference genes (glyceraldehyde-3-phosphate dehydrogenase -*GAPDH*, actin beta - *ACTB*, ribosomal protein L18 - *RPL18*, peptidylprolyl isomerase A *- PPIA*, tyrosine 3-monooxygenase/tryptophan 5-monooxygenase activation protein zeta - *YWHAZ*, ring finger protein 11 *- RNF11*, H3.3 Histone A *- H3F3A*, succinate dehydrogenase complex flavoprotein subunit A *– SDHA)* were analyzed using the geNorm function with the qbase+ package (Biogazelle, Zwijnaarde, Belgium) to identify the best reference genes^48^. Because it was more stably expressed (average geNorm M ≤ 0.5), *PPIA* was selected as the reference gene. A standard curve was included for each gene of interest as well as for the reference gene to confirm primer efficiencies for all targets were between 90% and 110%. The threshold cycle (Ct) for each sample was automatically calculated using the default settings within the SDS software (SDS 1.4, ABI).

The effect of the group on the ΔCt values for each gene was analysed by ANOVA with the animal treated as a random effect in a mixed model using the GLIMMIX procedure of SAS (SAS® OnDemand, SAS Institute, Cary, NC, USA). The model included the effect of the group, the heifer and the interactions between these terms. If the assumption of normality was not reached, according to the results of the Shapiro-Wilk test, ΔCt values were analyzed by the non-parametric Friedman test to determine the effect of the group when controlling for the heifer effect, using the FREQ procedure of SAS.

Target genes for qRT-PCR were selected either because of the fold-change difference observed in Experiment 1 (myosin light chain 4 – *MYL4*, interferon gamma *– IFNG*, cholinergic receptor muscarinic 3 – *CHRM3*, and proprotein convertase subtilisin/kexin type 2 *-PCSK2*), or through a supervised sparse Partial Least Square Discriminant Analysis (sPLS-DA; interleukin 1 receptor associated kinase 1 – *IRAK1*, CLK4 associating serine/arginine rich protein – *CLASRP*, negative elongation factor complex member B – *NELFB*, scribble planar cell polarity protein – *SCRIB*, RAB9A member RAS oncogene family – *RAB9A*, and sequestosome 1 - *SQSTM1*), applied with the mixOmics package^49^. Briefly, highly expressed genes (more than 100 counts per million in 6 or more samples) were evaluated for their ability to discriminate control and epidydimal sperm samples, reflected by the frequency that each gene correctly classified each sample after a leave-one-out cross-validation repeated 10 times.

## RESULTS

### Experiment 1: differential transcriptomic changes in the endometrium induced by epididymal and ejaculated sperm

Sequencing of endometrial explants exposed to epididymal sperm revealed a total of 1912 DEGs compared with control explants (FDR < 0.05; Supp. Table 2), of which 932 were upregulated and 980 downregulated (Fig. 2A). The 10 most up- and down-regulated DEGs are shown in Table 1. Two of the top GO biological processes associated with the upregulated genes were related to alpha-beta T cell activation (Fig. 3A). The TNF, NF-KB, and IL-17 signalling pathways were among the top KEGG pathways represented by the DEGs (Fig. 3B). Following this trend, “negative regulation NF-KB transcription factor activity” was among the top GO biological processes that were associated with the down-regulated genes (Fig. 3C). On the other hand, only three KEGG pathways were represented, including glycerophospholipid metabolism and vascular smooth muscle contraction (Fig. 3D).

**Table 1.**
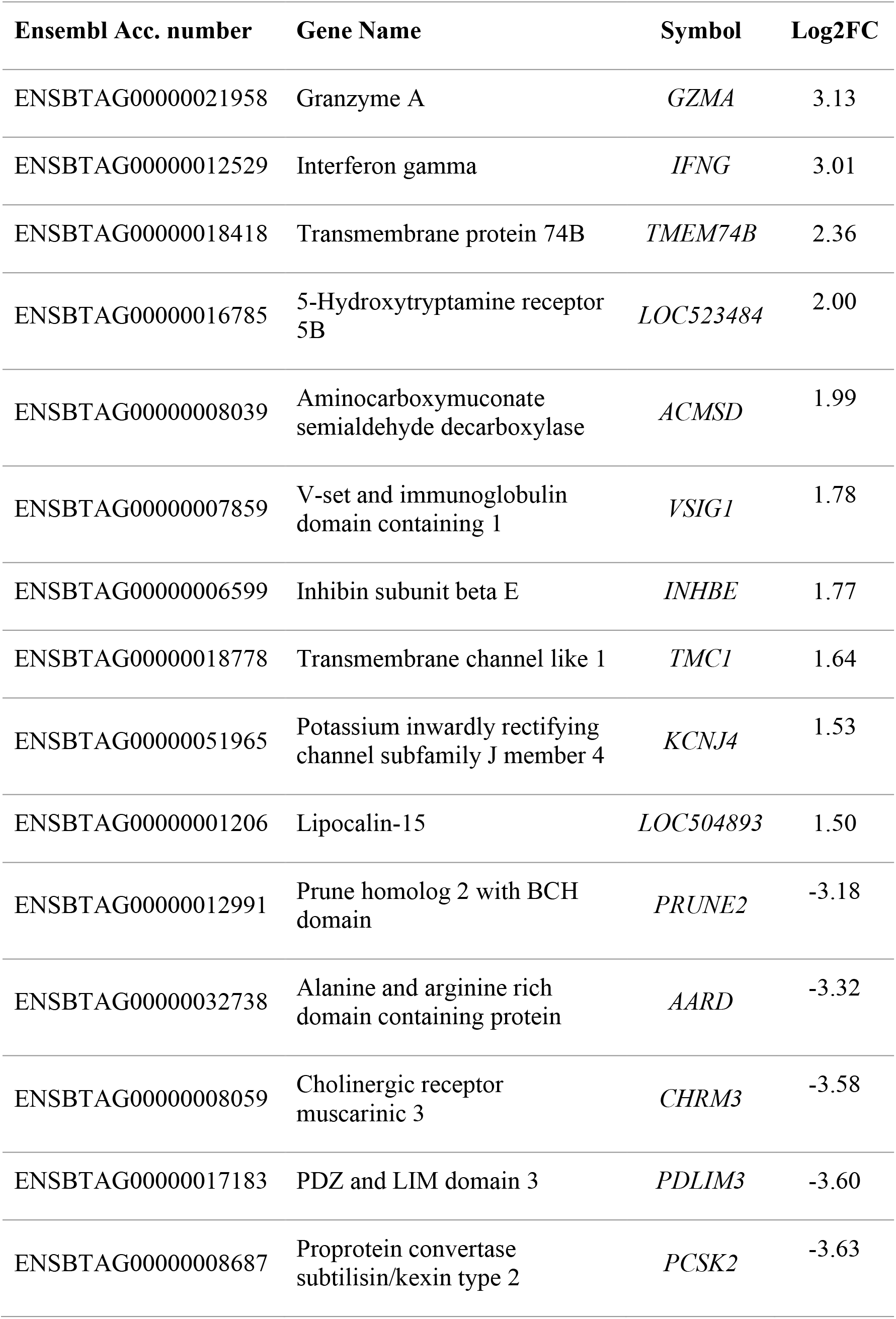
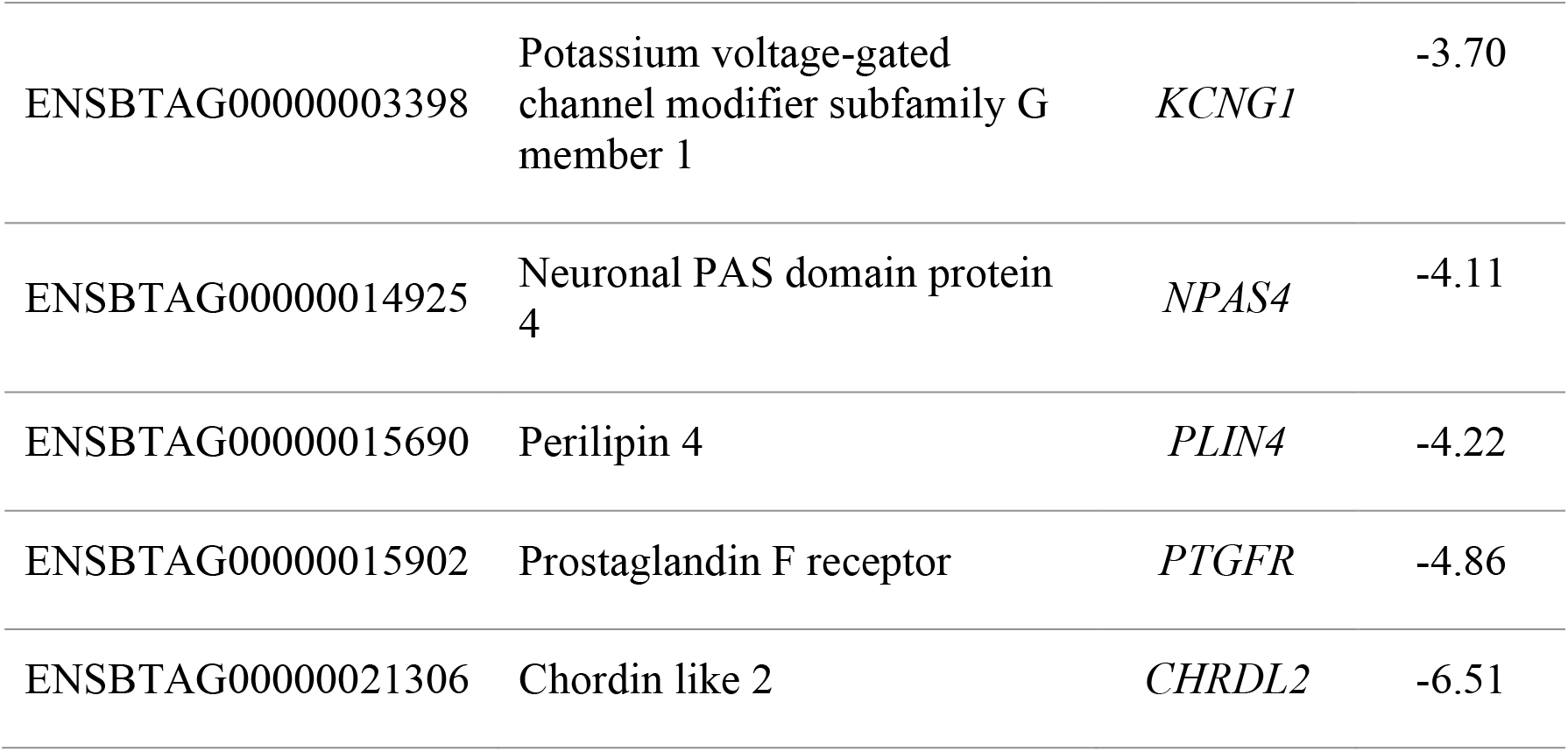
List of 10 most down- and up-regulated differentially expressed genes (FDR < 0.05) in bovine endometrial explants exposed to epididymal sperm compared with control explants.; Log2FC: logarithm with base 2 of fold change.

**Fig. 2.**
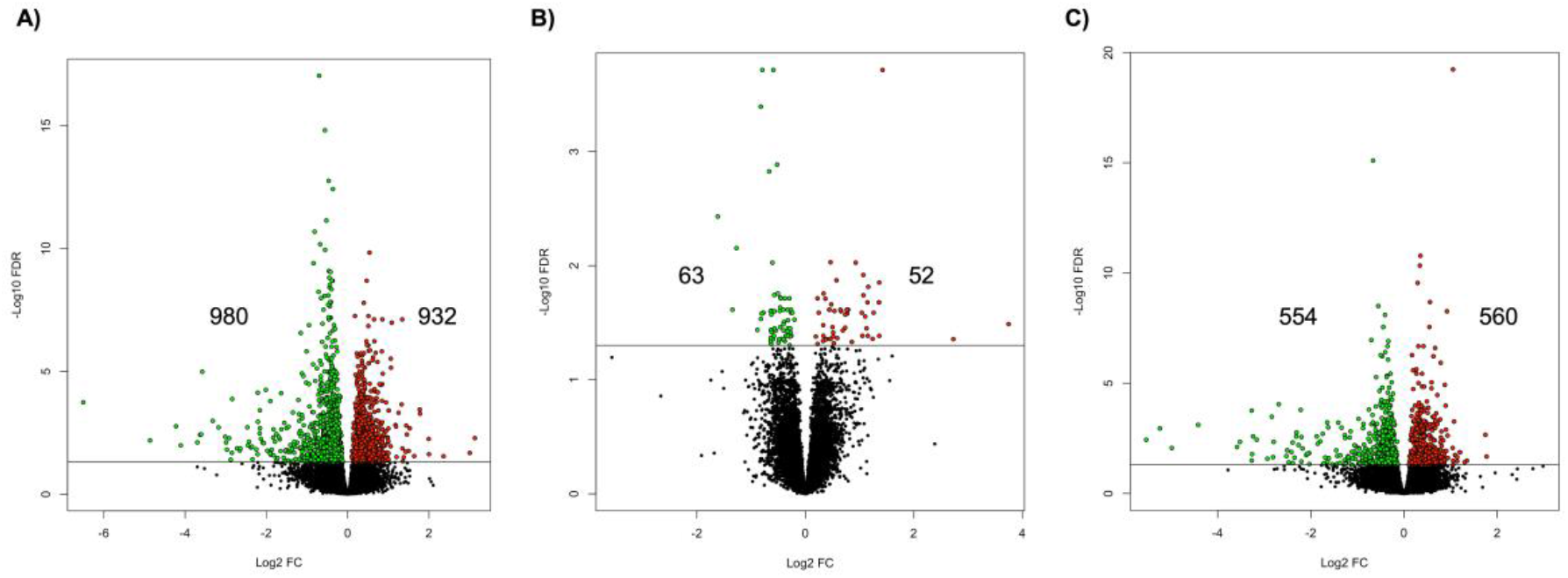
Volcano plot of differential expressed genes detected between: **A**) control explants and explants incubated with epididymal sperm, **B**) control explants and explants exposed to ejaculated sperm, and **C**) between explants incubated with epididymal and ejaculated sperm.

**Fig. 3.**
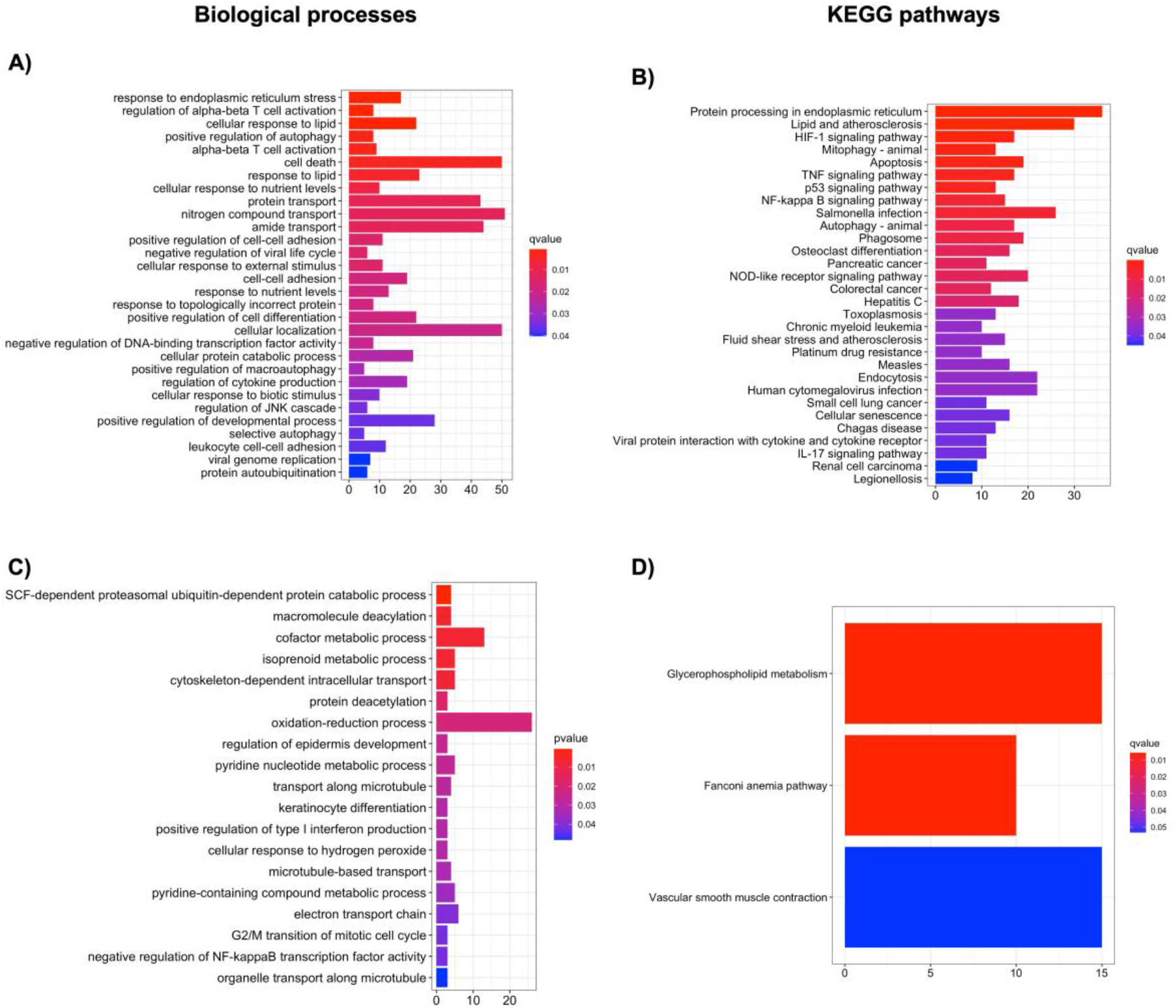
Top biological processes (**A** and **C**) and KEGG pathways (**B** and **D**) associated with the upregulated (**A-B**) and downregulated (**C-D**) endometrial genes induced by exposure to epididymal sperm

Exposure to ejaculated sperm, induced differential expression of 115 genes compared with control explants (52 up- and 63 downregulated; Fig. 2B), Table 2 contains the 10 most up- and down-regulated. Interestingly, only 36 DEGs were also differentially regulated in explants incubated with epididymal sperm vs. control explants (Supp. Table 3). Among the up- and downregulated genes, no enriched KEGG pathways were identified, nor GO biological processes among the downregulated genes. However, the top GO biological processes associated with the upregulated genes included “positive regulation of immunoglobulin production” and “regulation of acute inflammatory response” (Fig. 4).

**Table 2.** List of 10 most down- and up-regulated differentially expressed genes (FDR < 0.05) in bovine endometrial explants exposed to ejaculated sperm compared with control explants.; Log2FC: logarithm with base 2 of fold change.

| Ensembl Acc. number | Gene Name | Symbol | Log2FC |
| --- | --- | --- | --- |
| ENSBTAG000000007392 | Serpin peptidase inhibitor, clade A (alpha-1 antiproteinase, antitrypsin), member 14 | *SERPINA14* | 3.75 |
| ENSBTAG000000006354 | Haptoglobin | *HP* | 2.73 |
| ENSBTAG000000002414 | Potassium inwardly rectifying channel subfamily J member 10 | *KCNJ10* | 1.42 |
| ENSBTAG000000011833 | Glutamate ionotropic receptor AMPA type subunit 3 | *GRIA3* | 1.36 |
| ENSBTAG000000001021 | Cytochrome P450, subfamily I (aromatic compound-inducible), polypeptide 1 | *CYP1A1* | 1.36 |
| ENSBTAG000000008832 | C-C motif chemokine ligand 1 | *CCL1* | 1.36 |
| ENSBTAG000000011381 | Solute carrier family 30 member 3 | *SLC30A3* | 1.36 |
| ENSBTAG000000014921 | Interleukin 6 | *IL6* | 1.26 |
| ENSBTAG000000046010 | Basic helix-loop-helix family member a15 | *BHLHA15* | 1.24 |
| ENSBTAG000000053103 | Trichohyalin | *TCHH* | 1.16 |
| ENSBTAG000000021568 | Von Willebrand factor C and EGF domains | *VWCE* | -0.67 |
| ENSBTAG000000015651 |  |  | -0.77 |
| ENSBTAG000000024000 | Atonal bHLH transcription factor 8 | *ATOH8* | -0.79 |
| ENSBTAG000000046321 | TSPO associated protein 1 | *TSPOAP1* | -0.81 |
| ENSBTAG000000005685 | Hydroxysteroid 11-beta dehydrogenase 2 | *HSD11B2* | -0.82 |
| ENSBTAG000000004197 | UDP-GlcNAc:betaGal beta-1,3-N-acetylglucosaminyltransferase 4 | *B3GNT4* | -0.82 |
| ENSBTAG000000047810 | Coiled-coil domain containing 96 | *CCDC96* | -0.88 |
| ENSBTAG000000050871 | Mast cell protease 2 | *LOC100139881* | -1.27 |
| ENSBTAG000000048077 | Melanoma antigen family D, 4B | *MAGED4B* | -1.34 |
| ENSBTAG000000011579 | Cholinergic receptor muscarinic 1 | *CHRM1* | -1.61 |

**Fig. 4.**
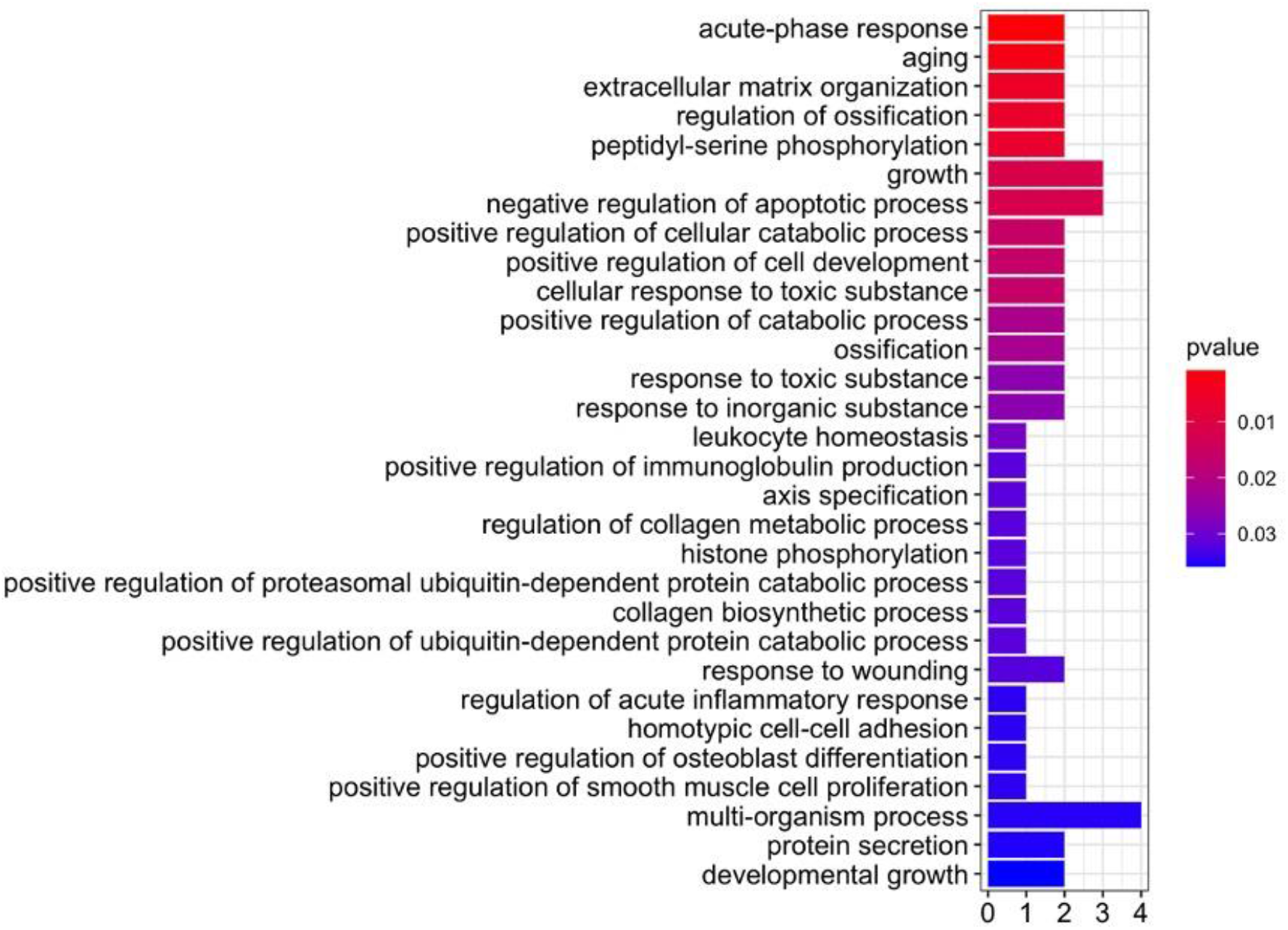
Biological process associated with the upregulated genes induced by incubation of endometrial explants with ejaculated sperm

When explants incubated with epididymal sperm were compared to explants exposed to ejaculated sperm, 1114 DEGs were found (554 up- and 560 downregulated; Fig. 2C). Regulation of T cells were, again, among the top biological processes associated with the upregulated DEGs, including “T cell mediated immunity” and “regulation of alpha-beta T cell activation”, as well as other immune related processes such as “response to interferon-gamma”, “lymphocyte differentiation” or “leukocyte activation” (Fig. 5A). Similarly, “negative regulation of NF-KB transcription factor activity”, “positive regulation of type I interferon production”, “regulation of cytokine-mediated signalling pathway” or “toll-like receptor signalling pathway” were among the top GO biological processes linked to the downregulated genes (Fig. 5C). On the other hand, “TGF-beta signalling pathway” or “Th17 cell differentiation” were some of the top KEGG pathways associated to the upregulated genes (Fig. 5B). Only three KEGG pathways were enriched by the downregulated DEGs, including “ribosome biogenesis” and “mRNA surveillance” pathways (Fig. 5D).

**Fig. 5.**
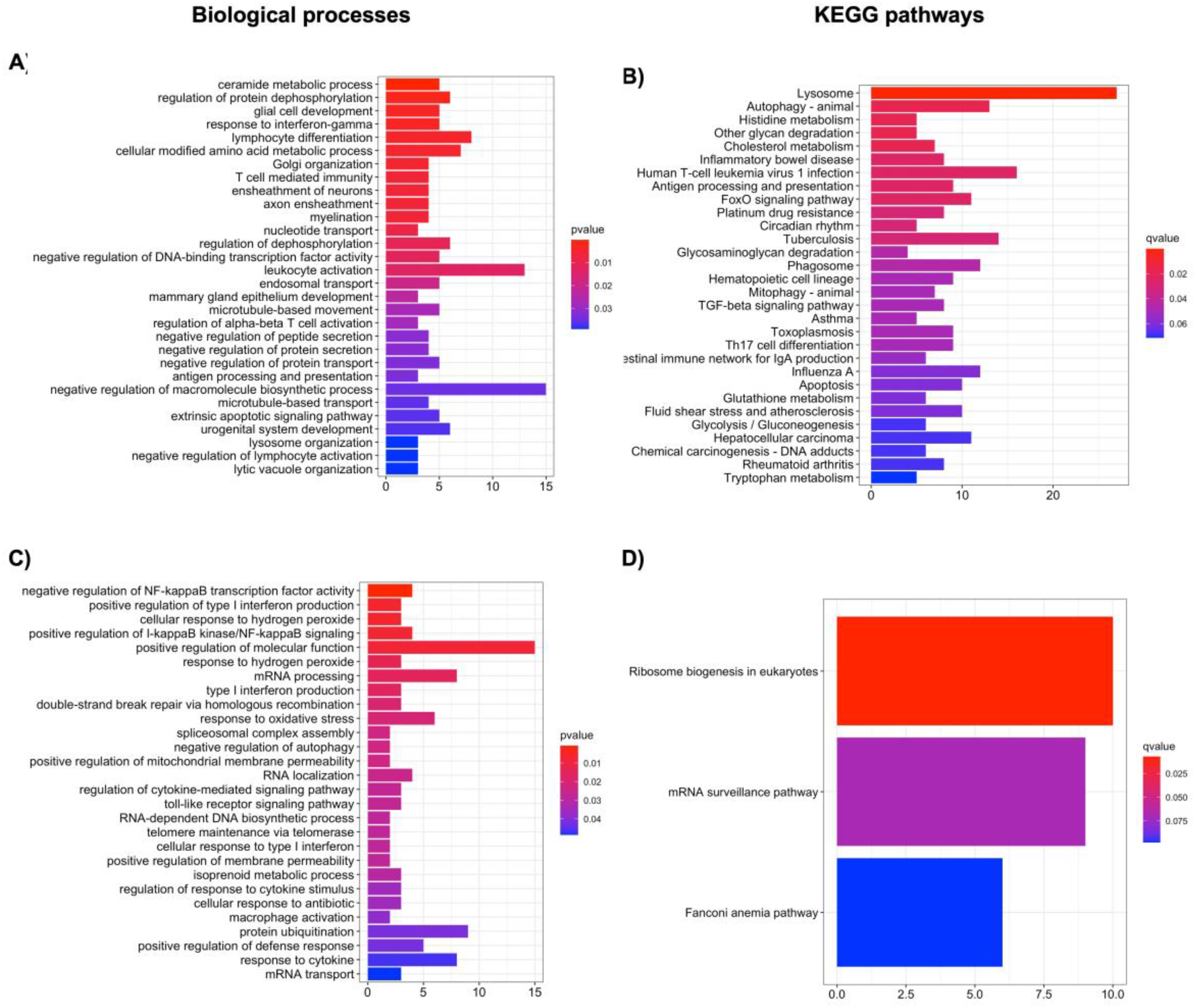
Top biological processes (**A** and **C**) and KEGG pathways (**B** and **D**) associated with the differentially upregulated (**A-B**) and downregulated (**C-D**) endometrial genes between the epididymal- and ejaculated sperm-exposed explants.

### Experiment 2: effect of SP on the ability of epididymal sperm to regulate endometrial gene expression

To determine whether SP factors were directly responsible for certain genes being differentially regulated by epididymal, but not ejaculated, sperm, the expression of a set of genes was analysed in endometrial explants exposed to epididymal sperm previously incubated, or not, with SP (Fig. 1B).

Incubation with SP inhibited the downregulation of *SQSTM1* in the endometrium induced by epididymal sperm, as evidenced by differences between Control and Epididymal (*P<* 0.01), but not Epididymal + SP groups (*P>* 0.05; Fig. 6C). A similar effect was observed in the upregulation of *MYL4* and *CHRM3*, and downregulation of *SCRIB*, although in this case, incubation with SP was not sufficient to completely block the effect of sperm exposure, as evidenced by differences not only between both sperm treatments (*P<* 0.05), but also between these and the control (*P<* 0.05; Fig. 6B).

**Fig. 6.**
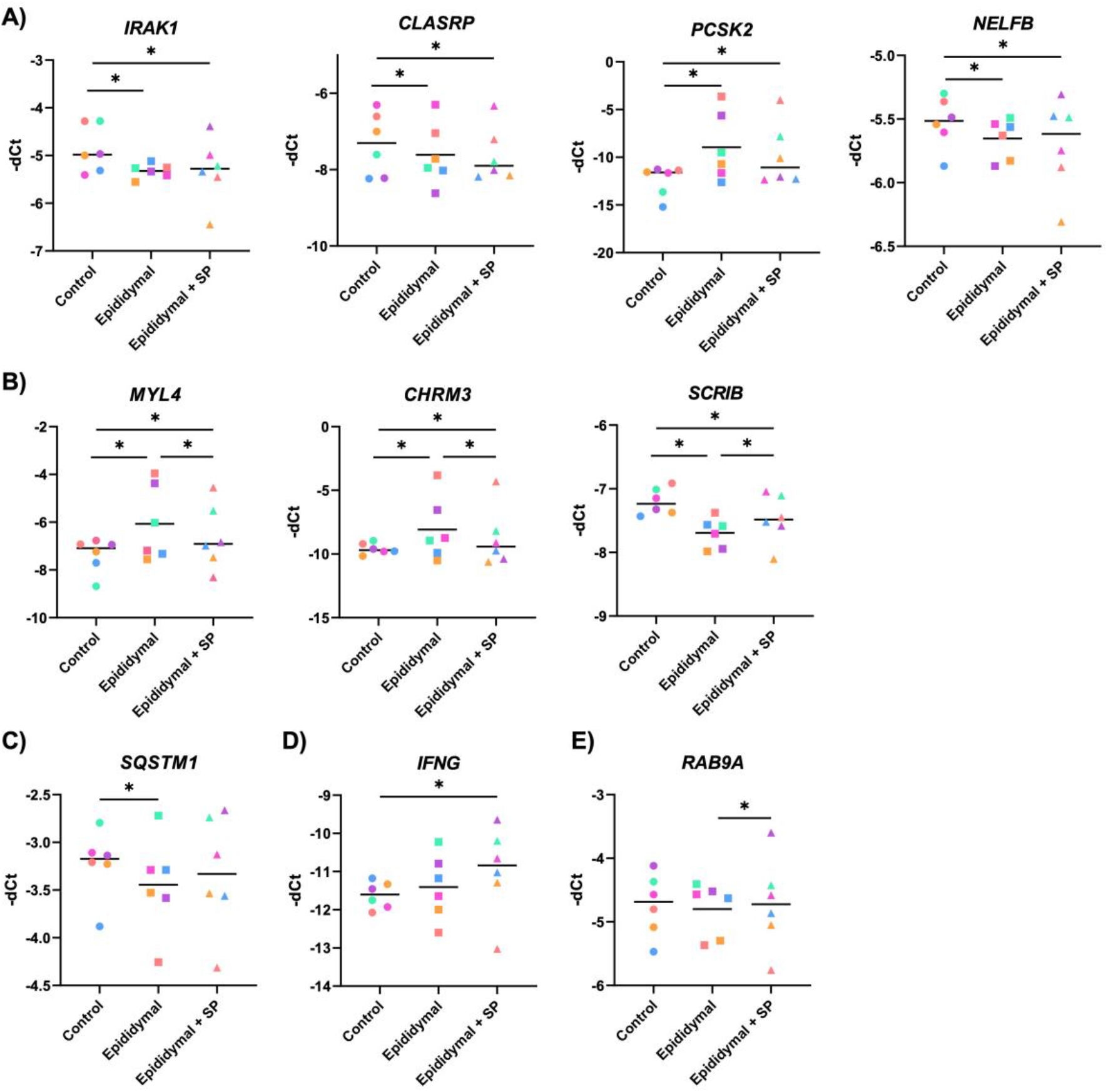
**A-E:** Dot plot representation (line as mean) of mRNA expression of *IRAK1, CLASRP, PCSK2, NELFB, MYL4, CHRM3, SCRIB, RAB9A, SQSTM1*, and *IFNG* as determined by q-rtPCR (-Delta Ct) in endometrial explants incubated with media alone (Control; n=6), epididymal sperm (Epididymal; n=6) or epididymal sperm previously exposed to SP (Epididymal + SP; n=6). Five expression patterns are observed in comparison with the control: **A)** regulated by epididymal sperm regardless of prior incubation with SP; **B)** differentially regulated by epididymal or epididymal + SP, **C)** regulated only by epididymal sperm, **D)** regulated only by epididymal + SP, and E) not regulated by epididymal sperm, regardless of prior incubation with SP. \**P*<0.05. Each colour represents explants recovered from the same heifer throughout all graphs.

On the other hand, *IRAK1, CLASRP, PCSK2* and *NELFB* expression differed between the control and both sperm treatments (Fig. 6A; *P<* 0.01), but a not between the Epididymal and the Epididymal + SP groups (*P>* 0.05). Interestingly, endometrial expression of *IFNG* was not affected by epididymal sperm unless previously incubated with SP (*P<* 0.05; Fig. 6D). Finally, *RAB9A* expression was not affected by epididymal sperm exposure, regardless of prior SP exposure or not, as no differences were observed between the control and the sperm treatments (*P>* 0.05; Fig. 6E).

## DISCUSSION

The main findings of the present study are: 1) both epididymal and ejaculated sperm induce changes in the endometrial transcriptome; 2) the endometrial response elicited by epididymal sperm is more dramatic (in terms of number of DEGs) than that of ejaculated sperm; and 3) addition of SP dampens the expression of certain endometrial genes by epididymal sperm.

We recently reported modest changes in the endometrial transcriptome 24 h after mating heifers to intact bulls (in which the ejaculate contains both sperm and SP), but not when heifers were mated to vasectomised bulls (in which the ejaculate lacks sperm and epididymal and testicular fluid^33^), suggesting that sperm-mediated factors were responsible for the observed effects. The present study extends these findings by evaluating whether these changes are elicited by factors secreted by the AGs and bound to sperm at ejaculation, or rather, by factors acquired during spermatogenesis and epididymal maturation. To this end, endometrial explants were co-incubated with cauda epididymal sperm (which have not been exposed to AG secretions) or ejaculated sperm.

Interestingly, the endometrial response elicited by epididymal sperm was more dramatic than that observed after incubation with ejaculated sperm, as evidenced by 1,912 genes found to be differentially regulated between control explants and explants incubated with epididymal sperm in comparison to 115 DEGs when control explants were compared to explants exposed to ejaculated sperm. In a recent study, incubation of mouse epididymal sperm with uterine epithelial cells for 16 h, induced an increased secretion of interleukin-6 (IL6), C-X-C motif chemokine ligand 2 (CXCL2), and colony stimulating factor 3 (CSF3), and a decrease in colony stimulating factor 2 (CSF2), TNF, LIF, and CCL2 production^50^. Of those cytokines, in our study, bull epididymal sperm only regulated expression of *CXCL2*, which was upregulated in comparison to control explants, whereas only ejaculated sperm induced upregulation of *IL6*. To the best of our knowledge, the study by Schjenken et al.^50^ is the only other study that has looked at the effect of epididymal sperm exposure on endometrial activity. However, bull epididymal-sperm-regulated pathways in the endometrium have been shown to be affected by exposure to ejaculated sperm in other species. Genes upregulated by epididymal sperm exposure were associated with T cell activation as well as IL17, TNF, and NF-KB signalling pathways. Similarly, in the mouse, mating to intact mice upregulated IL17 and NF-KB signalling, as well as Th1 and Th17 activation pathways, in comparison to mating to vasectomised males^50^. Enrichment analysis of endometrial DEGs induced by mating or AI in the sow, revealed an overrepresentation of pathways different to the ones observed in the bovine after incubation with either ejaculated or epididymal sperm, such as FoxO signalling, glycosaminoglycan biosynthesis or PI3K-Akt^51^. However, in a study focusing on the effect of mating on the regulation of glucocorticoid signalling through nuclear receptor subfamily 3 group C member 1 (NR3C1/GR) in the sow reproductive tract, nuclear receptor subfamily 3 group C member 1 (*NR3C1*), hydroxysteroid 11-beta dehydrogenase 2 (*HSD11B2*), FKBP Prolyl Isomerase 5 (*FKBP5*) and 4 (*FKBP4*), and prostaglandin-endoperoxide synthase (*PTGES1*) 1 and 2 (*PTGES2*) were differentially regulated by natural mating compared to cervical deposition of the sperm^52^. In the present study, epididymal sperm, but not ejaculated sperm, induced upregulation of *NR3C1, FKBP9* (another member of the NKBP family), and *PTGES3* in comparison to the control and upregulation of *HSD11B2* in comparison to ejaculated sperm, suggesting a similar role for epididymal sperm in the regulation of this pathway. Together, the data suggest that bull sperm, in the absence of factors derived from AG secretions can elicit changes in the endometrium similar to those induced by ejaculated sperm in other species. In fact, pathways related to T cell activation and mediated immunity, FoxO signalling, Th17 cell differentiation, or TGFB signalling were associated with genes upregulated by exposure to epididymal sperm in comparison to ejaculated sperm, while NF-KB signalling, type I interferon production or toll-like receptor signalling pathways were associated with genes downregulated by epididymal sperm in comparison to ejaculated sperm. Despite this, when comparing the DEGs induced by epididymal sperm in vitro to those regulated by mating to intact bulls in vivo^33^, only three common genes were found: C-X3-C Motif Chemokine Ligand 1 (*CX3CL1*), coagulation factor III (*F3*), and solute carrier family 24 member 2 (*SLC24A2*). Interestingly, CX3CL1 is a membrane-bound cytokine, regulated by cytokines such as IL1B, IFNG and TNFA, that promotes adhesion and chemotaxis of T lymphocytes, monocytes, natural killer (NK) cells, and dendritic cells (DCs)^53,54^. The sole receptor of CX3CL1 is CX3CR1, which is expressed in NK cells, monocytes and CD8+ T cells, and has also been shown to be present on sperm^55^. This suggest that sperm can both induce CX3CL1 expression and modulate its activity. Of note, in our in vivo study^33^, heifers were mated immediately after standing oestrus was observed and endometrial samples were collected 24 h later. In contrast, in the present study, endometrial explants were obtained 24 h after mating and subsequently exposed to sperm for 6 h. Considering that the LH peak occurs on average 9 h after standing oestrus^56^, at the time of mating, sperm in the in vivo experiment encountered a uterine environment that was subject to different hormonal concentrations to the one present when endometrial explants were recovered for the in vitro experiment. This could impact the regulation of sperm-endometrium interaction and explain, at least in part, the lack of more common DEGs found between this study and our prior in vivo one^33^. In addition, incubation time has also been shown to influence the response of bovine endometrial explants and epithelial cells to sperm^34,57^. Both time of sample collection and duration of incubation, could also be contributing factors to the differences in the regulation of gene expression observed: whereas *CX3CL1* was upregulated both by epididymal sperm exposure and mating to an intact bull, *F3* was upregulated by epididymal sperm but downregulated by mating to an intact bull, and *SLC24A2* was downregulated by epididymal sperm and upregulated by mating to an intact bull^33^. Differences in gene modulation might also be directly linked to differences in the ability of epididymal and ejaculated sperm to regulate the endometrial transcriptome. Indeed, although most of the 36 DEGs that were common to epididymal and ejaculated sperm behaved in a similar fashion (i.e., were similarly up-regulated or down-regulated in both groups), both forkhead box Q1 (*FOXQ1*) and Ras related glycolysis inhibitor and calcium channel regulator (*RRAD*) were upregulated by epididymal sperm but downregulated by ejaculated sperm. This suggests that modifications of sperm due to interaction with AG secretions modulate the subsequent response of the endometrium to sperm.

The more dramatic endometrial response elicited by sperm that had no contact with AG secretions, in comparison to ejaculated sperm, led to the hypothesis that factors of AG origin acquired at the time of ejaculation partially block the ability of sperm to interact with the endometrium. There is evidence in the literature which points to such an effect: bovine binder of sperm proteins, which are secreted by the seminal vesicles and make up approximately 50% of total bull SP protein^58^, are sialylated glycoproteins that bind to the sperm surface at ejaculation and partially lost after capacitation. The sialylated nature of these proteins is interesting considering that sialic acids, acquired during epididymal maturation and at ejaculation, have been hypothesised to play a role in masking potential antigenic sperm molecules until after sperm undergo capacitation^59–61^. In agreement with the hypothesis formulated, exposure to SP abrogated the downregulation of *SQSTM1* by epididymal sperm, and partially inhibited the upregulation of *MYL4* and *CHRM3* and downregulation of *SCRIB. SQSTM1* or *P62* encodes a multifunctional protein involved in autophagy and the regulation of the NF-KB signalling pathway^62^. In mice, uterine autophagy is high after mating, when the uterus exhibits an inflammatory response, and decreases around the time of implantation^63^. Considering that accumulation of P62 is associated with decreased autophagy^64^, it would seem that epididymal sperm stimulate uterine autophagy, whereas AG factors block this effect in cattle. Regarding *MYL4* and *CHRM3*, these genes participate in muscle contraction pathways; contractions propagate from the oviduct towards the cervix^65^ creating a flow of fluid that might be an important guiding mechanism for sperm due to their rheotaxis behaviour^66^. As evidence for the importance of these uterine movements for fertility, the number and intensity of uterine contractions positively correlates with pregnancy success in humans^67^. Although visual and olfactory stimuli from the bull are sufficient to induce small uterine contractions in cows^65^, the results reported herein point to differential regulation of this process by sperm and AG factors. This is also evident in the regulation of *SCRIB*, a gene which encodes a protein which plays a role in cell polarity, cell adhesion, proliferation and Hippo signalling^68^. In mice, *Scrib* is required for primary decidual zone formation and pregnancy success^69,70^, and there is also evidence for a similar effect in humans, where *SCRIB* plays a role in the changes of endometrial cell polarity required for decidualisation^71^. In contrast to all of these genes, downregulation of *IRAK1, CLASRP*, and *NELFB*, and upregulation of *PCSK2* was induced by epididymal sperm in the presence or absence of SP.

In conclusion, under the present experimental conditions, bull epididymal sperm regulated endometrial pathways which have also been observed to be regulated by exposure to sperm in the mouse and pig uterus. Factors of AG origin appear to modulate the interaction between sperm and the endometrium, affecting the expression of epididymal-sperm-regulated genes involved in muscle contraction, autophagy, and endometrial cell polarity, and inducing differential regulation of a different set of genes. However, the impact of the endometrial regulation driven by sperm and AG factors on cattle fertility remains to be elucidated.

## Acknowledgements

The authors wish to thank the National Cattle Breeding Centre for donating the fresh bull ejaculates used in this study, and Dr. John A. Browne (UCD) for his technical assistance.

## Author contributions

The study was designed by BFF, JMS and PL. BFF and PL secured funding. JMS and MM carried out the animal work. JMS, BFF and SBA carried out the laboratory work. TES performed RNA-sequencing, and MBR and SKB conducted bioinformatic analyses. BFF wrote the manuscript with extensive contribution from JMS, PL and MBR. All the authors read and approved the final manuscript.

