## Supplementary data for "Sperm exposure to accessory gland secretions alters the transcriptomic response of the endometrium in cattle"

**Supplementary Table 1**

Site and size of preovulatory follicle per animal used in the study.

|  | **Heifer** | **Site of preovulatory follicle (right/left ovary)** | **Size of preovulatory follicle** |
| --- | --- | --- | --- |
| **Experiment 1** | 1 | Right | Burst |
|  | 2 | Left | 14 x 11 mm |
|  | 3 | Left | 10 x 10 mm |
|  | 4 | Right | 15 x 12 mm |
|  | 5 | Left | 12 x 14 mm |
| **Experiments 2 and 3** | 1 | Left | 20 x 21 mm |
|  | 2 | Right | Burst |
|  | 3 | Left | 17 x 18 mm |
|  | 4 | Right | 12 x 16 mm |
|  | 5 | Right | 17 x 15 mm |
|  | 6 | Right | 19 x 17 mm |

**Supp Table 3.**

List of common endometrial differentially expressed genes (FDR < 0.05) induced by epididymal (Epid) or ejaculated (Ejac) sperm. Log2FC: logarithm with base 2 of fold change.

| Ensembl Acc. number | Symbol | Log2FC  Epid vs Control | Log2FC  Ejac vs Control |
| --- | --- | --- | --- |
| ENSBTAG00000032481 | *DAPL1* | 1.06 | 0.76 |
| ENSBTAG00000046773 | *MCOLN2* | 1.00 | 1.15 |
| ENSBTAG00000002414 | *KCNJ10* | 0.89 | 1.42 |
| ENSBTAG00000039992 | *TUBA1C* | 0.86 | 0.65 |
| ENSBTAG00000014465 | *SERPINE1* | 0.83 | 0.86 |
| ENSBTAG00000015127 | *SDC4* | 0.73 | 0.57 |
| ENSBTAG00000021594 | *TDRKH* | 0.66 | 0.79 |
| ENSBTAG00000052132 | *FOXQ1* | 0.65 | -0.40 |
| ENSBTAG00000040518 | *LOC112443216* | 0.65 | 0.51 |
| ENSBTAG00000004630 | *COMP* | 0.63 | 0.93 |
| ENSBTAG00000045588 | *LOC100298356* | 0.62 | 0.45 |
| ENSBTAG00000021686 | *MELK* | 0.62 | 0.66 |
| ENSBTAG00000013960 | *PSAT1* | 0.61 | 0.50 |
| ENSBTAG00000011661 | *CEP152* | 0.52 | 0.59 |
| ENSBTAG00000046314 | *TBC1D8B* | 0.47 | 0.48 |
| ENSBTAG00000010763 | *DUSP16* | 0.43 | 0.37 |
| ENSBTAG00000008849 | *SORT1* | 0.42 | 0.39 |
| ENSBTAG00000013009 | *AURKA* | 0.38 | 0.46 |
| ENSBTAG00000014727 | *RFC4* | 0.33 | 0.33 |
| ENSBTAG00000036262 | *GBE1* | 0.30 | 0.38 |
| ENSBTAG00000013929 | *RRAD* | 0.27 | -0.29 |
| ENSBTAG00000003354 | *SMCHD1* | 0.25 | 0.25 |
| ENSBTAG00000035226 | *TOR1AIP1* | 0.22 | 0.22 |
| ENSBTAG00000009293 | *EVL* | -0.23 | -0.27 |
| ENSBTAG00000016268 | *XRCC1* | -0.24 | -0.28 |
| ENSBTAG00000020040 | *LPCAT4* | -0.32 | -0.32 |
| ENSBTAG00000019530 | *KDM8* | -0.40 | -0.38 |
| ENSBTAG00000010138 | *SEMA3B* | -0.48 | -0.50 |
| ENSBTAG00000006404 | *CENPT* | -0.51 | -0.47 |
| ENSBTAG00000013523 | *OSBPL7* | -0.56 | -0.24 |
| ENSBTAG00000038132 | *AIFM3* | -0.58 | -0.57 |
| ENSBTAG00000021568 | *VWCE* | -0.66 | -0.67 |
| ENSBTAG00000004197 | *B3GNT4* | -0.79 | -0.82 |
| ENSBTAG00000046321 | *TSPOAP1* | -0.80 | -0.81 |
| ENSBTAG00000048486 | *IQANK1* | -0.84 | -0.47 |
| ENSBTAG00000048077 | *MAGED4B* | -1.64 | -1.34 |

**Supp Fig 1**

Principal component analysis (PCA) plot of **A)** raw data and **B and C)** data after the individual heifer effect was removed with the CombatSeq function from the sva packages. Colours in **A and B**  correspond to individual heifers (3 treatments/samples per heifer), whereas in graph **C**, colours correspond to different treatments.

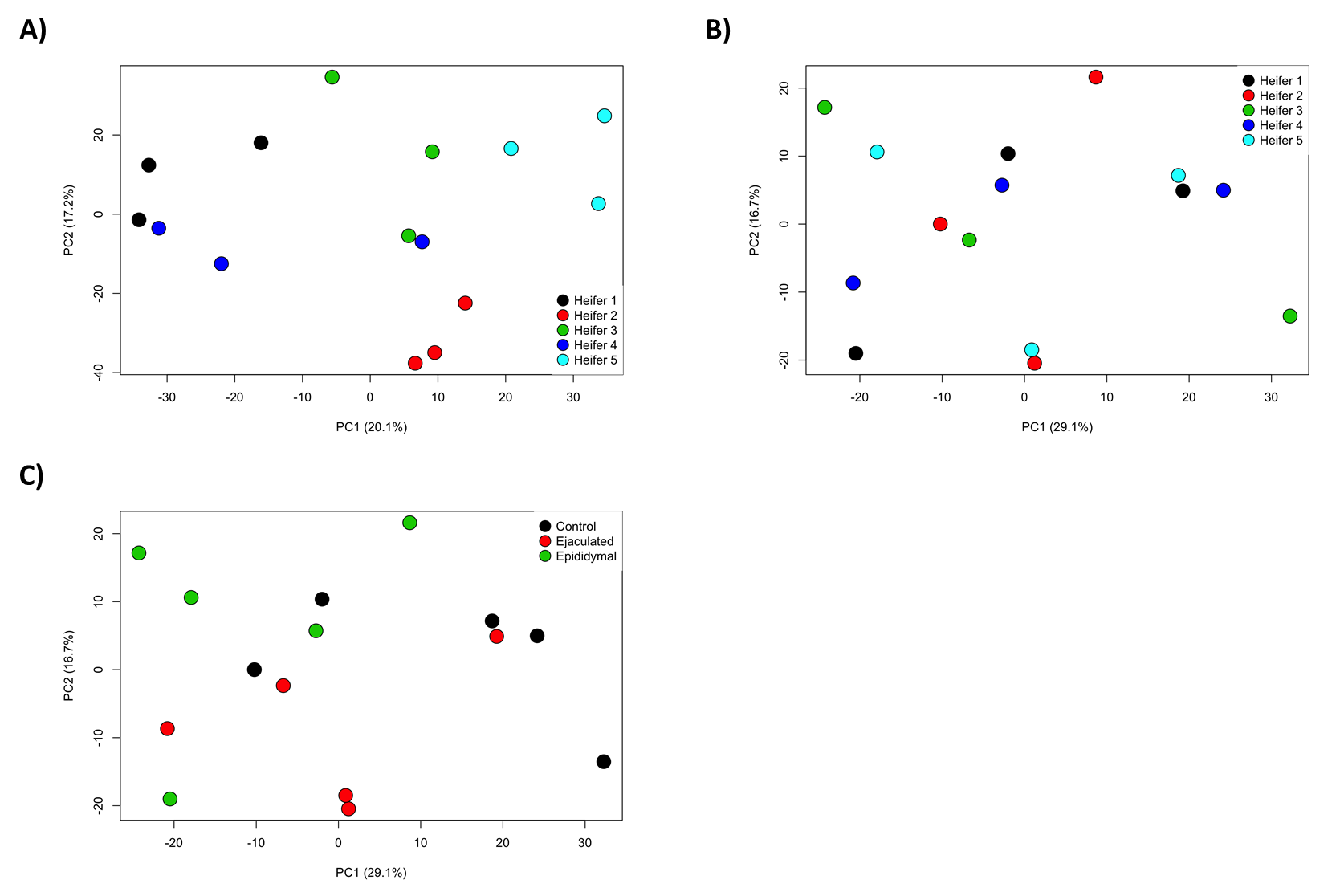
